# Maneuverable flight evolved with forked tails and opportunities for extrapair mating in swallows and martins (Aves: Hirundininae)

**DOI:** 10.1101/2023.12.02.569718

**Authors:** Masaru Hasegawa

**Affiliations:** Department of Environmental Science, Ishikawa Prefectural University, Nonoichi, Ishikawa 921-8836, Japan

**Keywords:** aerodynamic function, cost of ornamentation, flight cost, locomotor performance, plumage ornamentation

## Abstract

Whole-organism performance in relation to ornamentation is often examined to evaluate the cost of ornamentation, assuming that high performance is favored by viability selection. These studies typically conduct experimental manipulations of ornamentation, which potentially impair phenotypic integration with compensatory traits, making it difficult to clarify performance function of ornamentation. Here, we adopted an alternative approach, macroevolutionary analysis, and examined the flight performance of swallows (Aves: Hirundininae) in relation to tail fork depth to clarify evolutionary force favoring the ornamentation. We found that a measure of flight performance, presence of notable non-straight flight including maneuvering, turning, and swerving, peaked at intermediate fork depth, which appear to support viability selection for moderately forked tails. However, the quadratic relationship was found only in males, and hirundines with high opportunities for extrapair mating had higher probability of non-straight flight, indicating the importance of sexual selection. The current findings indicate that the flight performance of hirundines evolved through sexual selection, at least partially; thus, its relationship with forked tail might not clarify the viability cost of ornamentation. Whole-organism performance should be carefully interpreted when deducing the cost function, and thus, the evolutionary driver, of ornamentation.

---

Animals sometimes possess conspicuous ornaments, such as vivid coloration and weird appendages (Andersson 1994; Hill & McGraw 2006). Such ornamentation is often explained by sexual selection, i.e., reproductive advantage, for example, for attracting mates and repelling rivals (Anderson 1994). However, it is often difficult to clarify the evolutionary force behind ornamentation, because conspicuous traits can enhance viability in unexpected ways, for example, the colorful plumage of birds might startle prey, serve as a warning signal, enhance thermoregulation, provide acute vision, and so on (see Bortolotti 2006 for a review). Moreover, traits that have mainly evolved through viability selection can be a target of sexual selection as a byproduct, for example, due to high detectability or information content, and thus, current sexual function can be different from the past evolutionary forces behind ornamentation (i.e., exaptation; Bergstrom & Dugatkin 2016). Then, it is not surprising that researchers focus on the cost function of elaborate ornamentation, because ornaments that have evolved through sexual selection should accompany viability cost counterbalancing sexual selection, different from those evolved through viability selection, which by definition is close to the viability optimum (e.g., see Evans & Thomas 1997).

Out of the several cost types, performance cost is a major cost, particularly for structural ornamentation, such as long tails in birds, which can impair locomotor performance (e.g., a peacock’s train; Thavarajah et al. 2016). Empirical studies typically estimate performance costs through experimental manipulation of ornaments in a given population, by which possible confounding factors can be controlled. A representative example is that on forked tails in swallows (e.g., Buchanan & Evans 2000; Park et al. 2000; Matyjasiak et al. 2004, 2009). Deeply forked tails of swallows, particularly those of the barn swallow *Hirundo rustica,* are a classic example of sexually selected traits through social and extrapair mate choice (e.g., Møller 1988; reviewed in Møller 1994; Turner 2006; Romano et al. 2017), but might also have some aerodynamic function. Forked tails, i.e., triangular shaped tails when spread, can be regarded as a “delta wing,” improving the lift-to-drag ratio (until outermost tail feather length is twice as long as central tail feather length, beyond which a forked tail cannot acquire triangular shape at maximum spread, 120°; Thomas 1993). Even outermost tail feathers protruding beyond the maximum continuous span (i.e., “streamer” part) can provide additional lift, which can reduce turn radius (i.e., improving maneuverability; Norberg 1994). In support of these theories, when experimentally manipulating outermost tail feathers in the barn swallow, Evans and colleagues found that estimated peak aerodynamic performance located only around 10 mm shorter from the current outermost tail feathers, suggesting that most forked tails evolved through viability selection for aerodynamic function (e.g., Evans 1998; Buchanan & Evans 2000; Rowe et al. 2001). However, their studies, like those of many others that manipulated ornaments, did not take into account the presence of compensatory traits (Hasegawa 2023; also see Oufiero & Garland 2007; Husak & Swallow 2011 for general cautions of ignoring compensatory traits). According to Norberg (1994, p.231), *any experimental shortening and lengthening of the outer tail feathers is likely to upset an original co-adapted character set.* In other words, reducing forked tails might deteriorate phenotypic integration with compensatory traits (e.g., wing size and shape; Møller 1996), and thus, impair whole-organism performance (Fig. 1, light blue line; Hasegawa 2023). Therefore, these experiments were not conclusive, and the evolutionary force favoring forked tails remains controversial (e.g., Barbosa 1999; Møller & Barbosa 2001; Matyjyasiak et al. 2009 for early disputes; Hasegawa & Arai 2020, 2021, 2022 for recent discussions).

**Fig. 1.**
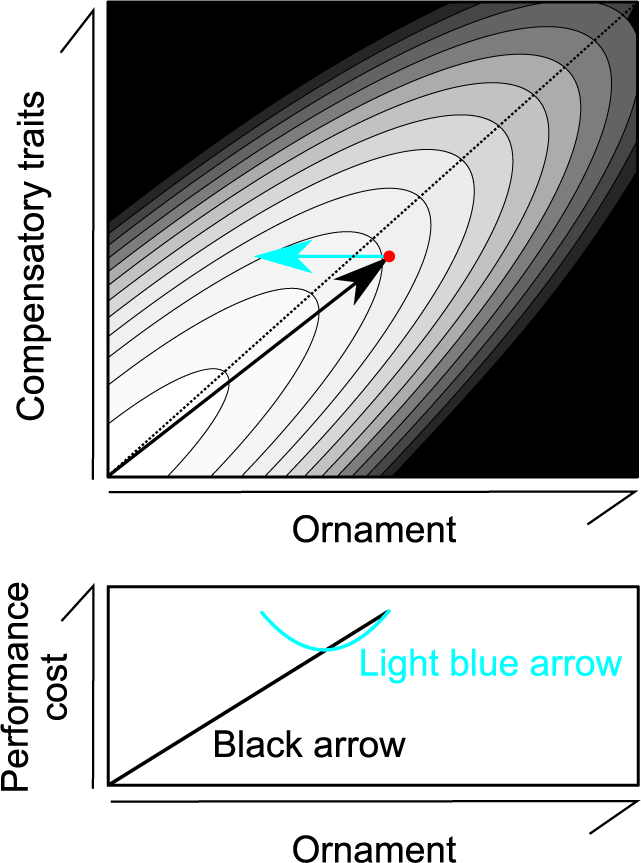
A hypothetical performance surface in relation to ornament and phenotypic expression of compensatory traits (upper panel). Darker background coloration indicates lower performance (note that the best performance is attained in the lower left corner). Black arrow indicates hypothetical evolutionary pathway to the current state (filled circle). Light blue arrow indicates phenotypic change when ornamentation was experimentally reduced. Lower panel shows performance cost measurement, denoted as best performance minus performance of the focal coordinate, in relation to ornament expression, with black or light blue arrows. The evolutionary pathway to the current state is biased toward the right because intense sexual selection favored long tails. Dotted line is shown as a ridge line. See Hasegawa (2023) for detailed explanation. Illustration was given by Hasegawa (2023).

An alternative approach to study performance cost of ornamentation is a phylogenetic comparative analysis, in which the macroevolution of animal performance is examined in relation to ornamentation (e.g., Oufiero & Garland 2007; Oufiero et al. 2014). Assuming evolutionary continuity of ornamentation on an adaptive landscape (e.g., Matyjasiak et al. 2000, 2004, 2009; also see Arnold 2023 for a review on theory), we can study how performance changed with the evolution of ornamentation without impairing its phenotypic integration with compensatory traits (Fig. 1, black line). Because hirundines are hyperaerial insectivores foraging in open space, there are abundant descriptions on their flight performance (Turner & Rose 1994), providing a suitable opportunity to study performance changes in relation to the evolution of ornamentation. A series of recent macroevolutionary studies of this clade (e.g., Hasegawa & Aria 2020, 2021, 2022; Hasegawa et al. 2022), together with detailed within-species studies in hirundines (e.g., Møller 1994; Magrath & Elgar 1997; Turner 2006), provide useful information on sexual selection and foraging ability as well.

Here, using phylogenetic comparative analyses of swallows and martins (Aves: Hirundininae), we examined the evolution of flight performance in relation to the expression of forked tails. We predicted that, if sexual selection favors deeply forked tails, flight performance would decrease linearly with fork depth. This prediction contrasts with that of an alternative hypothesis in which viability selection favors forked tails due to aerodynamic function. Under this hypothesis, we predicted that flight performance, notably maneuverable flight, should be enhanced by forked tails, at least to some points (see above). Because tail fork depth covaries with indices of sexual selection (Hasegawa & Arai 2021, 2022), we also analyzed these potentially confounding factors. Finally, based on the assumption that high aerodynamic performance improves foraging ability (e.g., Norberg 1994; Rowe et al. 2001), we tested whether foraging ability increases with our measures of flight performance.

## METHODS

### Data collection

Information on morphology (wing length, as a measure of body size, and relative fork depth), prey size (large or small), and migratory habits (migrant or not) of hirundines were obtained from Turner & Rose (1994), as described before (e.g., Hasegawa & Arai 2020, 2021, 2022; Hasegawa et al. 2022). Relative fork depth was calculated as (fork depth)/(central tail feather length), where fork depth is the difference between outermost and central tail feather lengths. In hirundines, foraging on large prey is thought to be more efficient than foraging on small prey (e.g., Turner 1982), and thus, a dichotomous prey category (large vs. small) provides useful information on foraging ability (Hasegawa & Arai 2022). Information on sexual plumage dimorphism, including all types of notable plumage characteristics in the wild, and incubation type as indices of sexual selection and opportunities for extrapair mating, respectively, originally obtained from Turner & Rose (1994), were given elsewhere together with the rationales (Hasegawa & Arai 2022). Incubation type is notable, because it explains tail fork depth and its sexual dimorphism (Hasegawa & Arai 2020, 2022) and sperm size (Hasegawa et al. 2022). Furthermore, species with female-only incubation exhibited three times higher extrapair paternity rate than those with biparental incubation (Hasegawa & Arai 2020), demonstrating the validity as an index of opportunities for extrapair paternity, as predicted by a temporal tradeoff between extrapair mating effort and incubation effort (also see Magrath & Elgar 1997 for within-population relationship between male incubation and paternity).

In addition, we obtained information on flight performance using the same literature, mainly in the “food and behavior” section (or “foraging and food” in some cases; Turner & Rose 1994). Although our focus here is non-straight flight, including maneuvering, turning, and swerving, which is the key theoretical focus (see above), we also collected information of gliding flight and fast flight, as additional measures of flight performance that are potentially associated with tail fork depth (e.g., Cuervo et al. 1996; Rowe et al. 2001). Concerning non-straight flight, a binary variable (exhibiting or not exhibiting notable non-straight flight), we designated a species as a non-straight flyer when there was “turning,” “maneuvering,” “swerving,” and similar terms in Turner & Rose (1994). Because these terms were mainly used for fast flyers, we also included non-straight flight if “zigzagging,” “erratic,” and similar wording described non-straight flight. Considering gliding flight, a binary variable (gliding or not), we designated species as having gliding flight when the focal species were described to have “gliding” flight in Turner & Rose (1994), although we excluded species with few gliding or similar expression from this category. Concerning fast flight, a binary variable (fast or not), we regarded species as having fast flight, when “fast,” “swift,” “rapid,” or similar wording were used in Turner & Rose (1994). The dataset for the current study can be found in Table S1.

### Statistics

A Bayesian phylogenetic mixed model with binomial error distribution, was used to examine each measure of flight performance (i.e., non-straight flight, gliding flight, and fast flight). To account for potentially confounding variables (i.e., body size, migratory habits, here), we included these variables as covariates. To account for phylogenetic uncertainty, we fit models with each tree and applied multimodel inference using 1,000 alternative trees obtained from birdtree.org (Garamszegi & Mundry 2014; Rubolini et al. 2015; see Sheldon et al. 2005 for an original phylogeny in swallows). As recommended for categorical variables, residual variance was fixed to one (de Villemereuil et al. 2013). We used a Gelman prior for the fixed effects while standardizing each continuous variable. Chains were run for 100,000 iterations, with a burn-in period of 60,000 and a thinning interval of 40 for each tree. Mean coefficient, 95% credible interval (CI), MCMC-based *P*-values (*P*_MCMC_), and phylogenetic signals were denoted. All analyses were conducted using the MCMCglmm function in the MCMCglmm package (Hadfield 2010) on the R platform (R Core Team 2021).

We used a discrete module in BayesTraits (Pagel & Meade 2006) to examine evolutionary transitions among states with the presence or absence of non-straight flight and incubation type (biparental vs. female-only incubation). Again, 1,000 trees were used to account for phylogenetic uncertainty (see above). Here, we ran 1,010,000 iterations with a burn-in period of 10,000 and a thinning interval of 1,000 (note that the phylogenetic tree was randomly chosen from 1,000 trees in each iteration). Means were denoted as the representatives of each transition rate. I also showed the posterior probability that the differences in transition rates would be higher (or lower) than zero (as *P*_MCMC_ values). The reproducibility of the MCMC simulation was assessed by calculating the Brooks–Gelman– Rubin statistic (Rhat), which must be <1.2 for all parameters (Kass et al. 1998), by repeating the analyses three times. Bayes factor was calculated by comparing the marginal likelihood of a dependent model that assumed the correlated evolution of sexual dimorphism and biparental incubation to that of an independent model that assumed the two traits evolved independently, using the stepping stone sampler implemented in BayesTraits. Bayes factor >2, 5–10, and >10 indicates positive, strong, and very strong evidence of correlated evolution, respectively (Kass & Raftery 1995; BayesTraits Manual Ver. 3). As before (e.g., Hasegawa & Arai 2022), statistical results supporting correlated evolution (see “Results” section) represented replicated co-distributions and bursts throughout hirundines, and thus, false positives due to a few influential evolutionary events would be unlikely (Maddison & FitzJohn 2015; Revell & Harmon 2022).

## RESULTS

### Relationship between flight performance and forked tail

Of the three measures of flight performance, gliding flight was not significantly explained by the depth of male fork tails (Table S2). This was also the case when we used female morphologies, instead of male morphologies (Table S2). Likewise, fast flight was not significantly explained by male or female tail fork depth (Table S3). Ancestral state reconstruction of the two measures of flight performance indicates that they were repeatedly evolved (Fig. S1).

Non-straight flights, including maneuverable flight, turning, and swerving, were repeatedly evolved (Fig. 2) and was significantly explained by the quadratic term of tail fork depth in males (Table 1): non-straight flight was most likely to be found in hirundines with intermediately forked tails (Fig. 3). When back-transformed from Table 1 (in which forked tails were standardized), the estimated depth of male fork tails that provided peak performance was 0.86 [95%CI = 0.55, 1.20, P_MCMC_ < 0.01]. When female tail fork depth instead of male one was used, neither quadratic nor linear term was significant (Table 1).

**Fig. 2.**
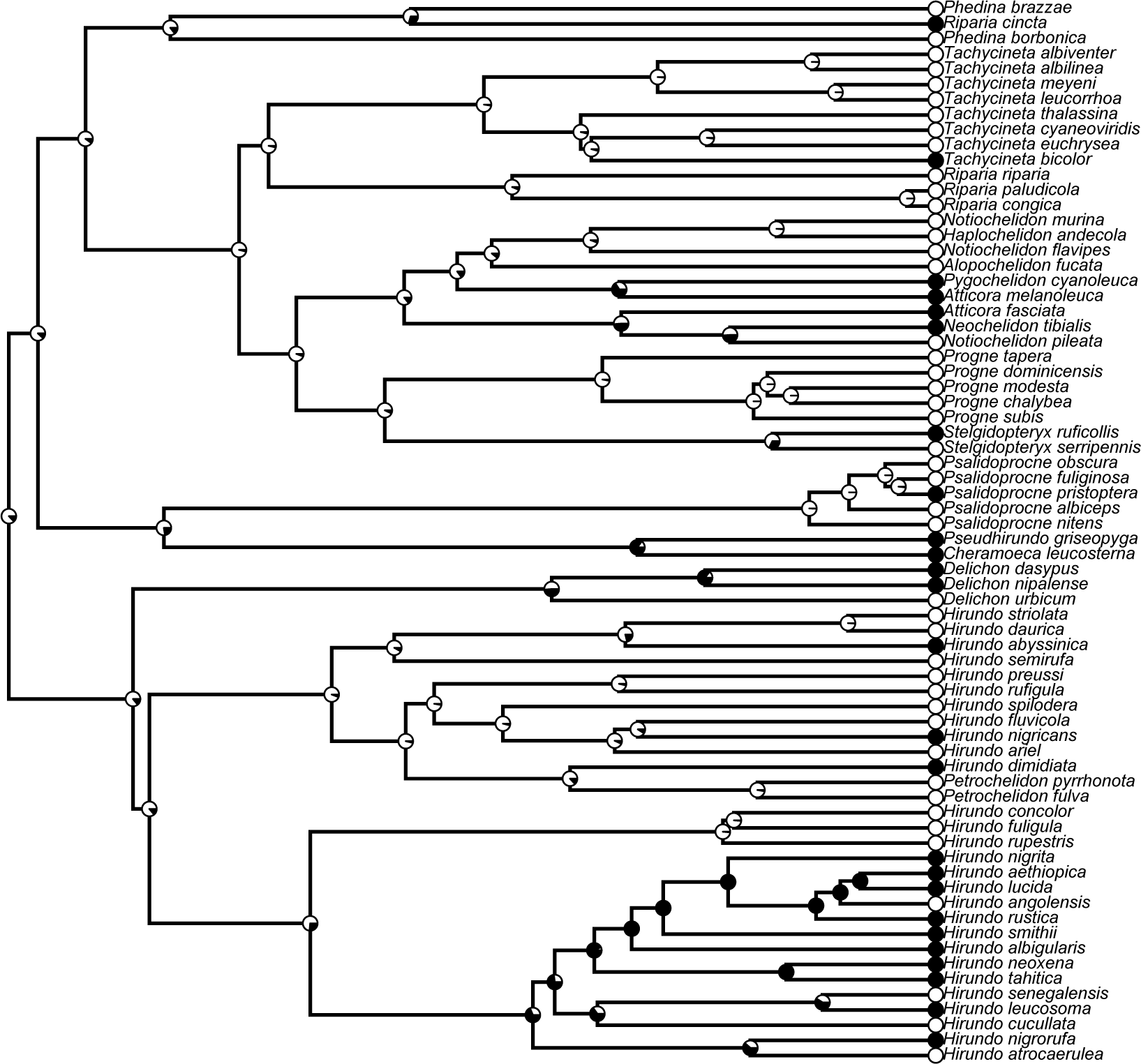
An example of ancestral character reconstruction of non-straight flight in swallows and martins (Aves: Hirundininae). The presence and absence of non-straight flight are indicated with black and white circles, respectively. The proportions of black and white at the nodes indicate the probability of the ancestral state. Here, I used the “ace” function in the R package “ape” (with model = “ER,” i.e., an equal-rates model; Paradis et al. 2005) and “plotTree” in the R package “phytools” (Revell 2012) for reconstructing ancestral character.

**Fig. 3.**
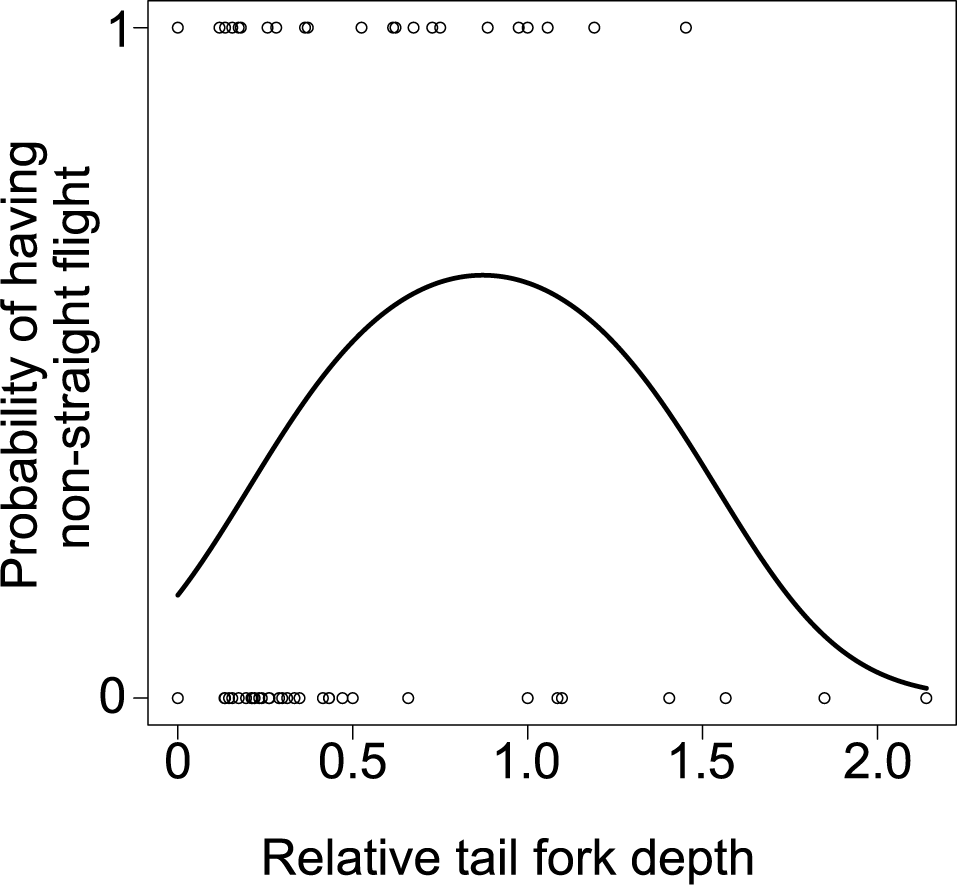
Relationship between non-straight flight and male tail fork depth in hirundines (n = 70). A simple logistic regression line is denoted for illustrative purposes. See Table 1 for formal analysis.

**Table 1.** Multivariable Bayesian phylogenetic mixed model with binary error distribution predicting the presence or absence of non-straight flight in relation to tail fork depth in male and female hirundines (both: n = 70).

| Variables | Estimates | 95% CI | $P_{\text{MCMC}}$ |
| --- | --- | --- | --- |
| Males |  |  |  |
| log(wing length) | -0.83 | -1.69, -0.02 | 0.03 |
| Migratory habit | 0.93 | -0.40, 2.31 | 0.16 |
| Tail fork depth | 1.27 | 0.24, 2.33 | 0.01 |
| (Tail fork depth)<sup>2</sup> | -0.79 | -1.45, -0.20 | <0.01 |
| Phylogenetic signal = 0.64 |  |  |  |
| Females† |  |  |  |
| log(wing length) | -0.83 | -1.67, -0.03 | 0.03 |
| Migratory habit | 0.75 | -0.52, 2.09 | 0.25 |
| Tail fork depth | 0.84 | -0.05, 1.78 | 0.06 |
| (Tail fork depth)<sup>2</sup> | -0.24 | -0.93, 0.44 | 0.48 |
| Phylogenetic signal = 0.63 |  |  |  |

log(wing length) and tail fork depth were standardized to zero mean and unit variance before analysis.
†When removing the quadratic term of female tail fork depth, the estimate of the linear term became 0.64 (95% CI = -0.04, 1.35), $P_{\text{MCMC}} = 0.06$ .

To study the potential confounding factors of male fork depth, we examined non-straight flight in relation to incubation type as an index of opportunities for extrapair mating (see Methods). Incubation type significantly explained non-straight flight (Table 2): hirundines with female-only incubation (i.e., those with frequent opportunities for extrapair mating) had a higher probability of non-straight flight than those with biparental incubation (i.e., those with limited opportunities for extrapair paternity). When replacing incubation type with another index of sexual selection, sexual dimorphism in overall plumage characteristics, we found that this variable was far from significant (*P*_MCMC_ = 0.92; details not shown).

**Table 2.** Multivariable Bayesian phylogenetic mixed model with binary error distribution predicting the presence or absence of non-straight flight in relation to incubation type as a measure of opportunity of extrapair paternity in male hirundines (both: n = 41).

| Variables | Estimates | 95% CI | P<sub>MCMC</sub> |
| --- | --- | --- | --- |
| log(wing length) | -1.39 | -2.54, -0.28 | <0.01 |
| Migratory habit | 0.21 | -1.52, 2.01 | 0.83 |
| Tail fork depth | 0.80 | -0.58, 2.21 | 0.25 |
| (Tail fork depth)<sup>2</sup> | -0.58 | -1.37, 0.15 | 0.10 |
| Incubation type | -2.53 | -4.51, -0.60 | <0.01 |
| Phylogenetic signal = 0.50 |  |  |  |

log(wing length) and tail fork depth were standardized to zero mean and unit variance before analysis.

When incubation type was included into models of other measures of flight performance (i.e., gliding and fast flight), gliding flight was not significantly explained by incubation type (Table S4). However, fast flight was significantly explained by incubation type, in which hirundines with female-only incubation had a higher probability of fast flight (Table S4). When sexual dimorphism in overall plumage characteristics was used instead of incubation type, this variable was non-significant in the models of the two measures of flight performance (*P*_MCMC_ > 0.08; details not shown).

Finally, we examined the three measures of flight performance in relation to prey size, which is thought to be linked to foraging ability in hirundines, with those foraging on large prey having higher foraging ability than others (see Methods). Prey size was not significantly explained by the three measures of flight performance (Table 3). Excluding covariates did not change the results qualitatively (*P*_MCMC_ > 0.43; details not shown).

**Table 3.** Multivariable Bayesian phylogenetic mixed model with binary error distribution predicting prey size (large vs. small) in relation to measures of flight performance in male hirundines (both: n = 58).

| Variables | Estimates | 95% CI | P<sub>MCMC</sub> |
| --- | --- | --- | --- |
| log(wing length) | 1.41 | 0.49, 2.40 | 0.001 |
| Migratory habit | 0.37 | -0.96, 1.71 | 0.58 |
| Tail fork depth | -1.02 | -1.89, -0.17 | 0.01 |
| Gliding flight | -0.86 | -2.18, 0.45 | 0.19 |
| Fast flight | 1.00 | -0.40, 2.39 | 0.15 |
| Non-straight flight | 0.38 | -1.11, 1.91 | 0.63 |
| Phylogenetic signal = 0.44 |  |  |  |

log(wing length) and tail fork depth were standardized to zero mean and unit variance before analysis. The quadratic term of tail fork depth, once included in the model, was far from significant ( $P_{\text{MCMC}} = 0.75$ ).

### Evolutionary transitions

When examining evolutionary transitions between non-straight flight and incubation type, we found that a dependent model that assumed a correlated evolution of them better fit the data compared with an independent model that assumes the two traits evolved independently (Bayes factor = 2.74), indicating positive evidence for correlated evolution. In the dependent model, transition rate from the state with non-straight flight and biparental incubation was higher than the reverse transition (*P*_MCMC_ = 0.01; Fig. 4).

**Fig. 4.**
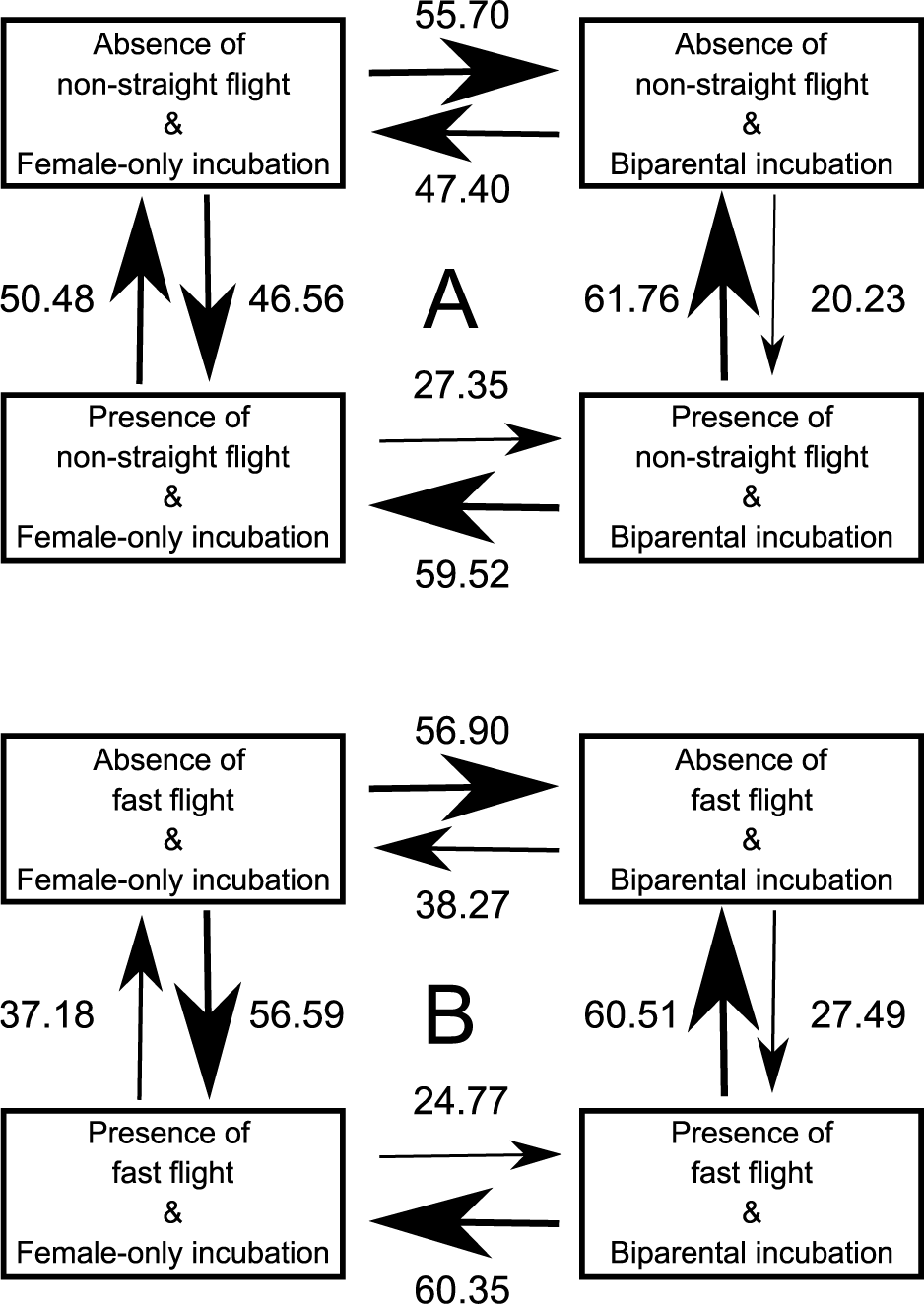
The most likely evolutionary transitions: (A) between states with presence or absence of non-straight flight and incubation type (female-only vs. biparental); and (B) between states with presence or absence of fast flight and incubation type (female-only vs. biparental) in swallows and martins (Aves: Hirundininae). Numbers next to arrows, reflected by arrow size, indicate transition rates between pairs of states.

When we used gliding flight instead of non-straight flight, the dependent model did not fit the data better than the independent model (Bayes factor = −0.11; note that negative values can be estimated). When we focused on fast flight; however, the dependent model fit the data better than the independent model (Bayes factor = 5.52), indicating strong evidence for correlated evolution between incubation type and fast flight. In the dependent model, transition rate from the state with fast flight and biparental incubation was higher than the reverse transition (*P*_MCMC_ = 0.022; Fig. 4).

When examining evolutionary transitions of prey size in relation to non-straight flight, a dependent model that assumed their correlated evolution did not fit the data better than the independent model (Bayes factor = −0.87). This was also the case when we examined prey size in relation to gliding flight (Bayes factor = −0.59) and fast flight (Bayes factor = 0.72).

## DISCUSSION

The main finding of the current study is that the presence of notable non-straight flight of swallows was explained by the quadratic term of male tail fork depth: hirundines with intermediate fork depth show the highest probability of having non-straight flight, as expected by aerodynamic function of forked tails (Thomas 1993; Norberg 1994). The estimated “optimum” fork depth for non-straight flight (relative fork depth = 0.83; see Results) is somewhat shallower than that reported by previous manipulative experiments (e.g., 1.21 for Buchanan & Evans 2000, based on the relative fork depth of the barn swallow), as predicted by the presence of compensatory traits (Fig. 1; also see fig. S2 in Hasegawa 2023). However, contrary to our expectation, flight performance was explained by a correlate of male tail fork depth, opportunities for extrapair mating, indicating the confounding effect of sexual selection. This finding is further reinforced by the observation that male, and not female, fork depth explained non-straight flight. We discuss the observed pattern in ecological and evolutionary perspectives.

Incubation type, i.e., an index of opportunities for extrapair mating, was related to two measures of flight performance (i.e., non-straight flight and fast flight) after tail fork depth was statistically controlled (Tables 2 & S3), indicating that these relationships were partially independent from tail fork depth. In addition, we found that an index of total sexual selection, sexual dimorphism in overall plumage characteristics, did not explain flight performance, indicating the importance of opportunities for extrapair mating rather than sexual selection in general. An alternative explanation that male incubation behavior itself (i.e., whether males participate in incubation) affects flight performance is unlikely, because incubation participation does not need particular flight performance (note that male hirundines typically participate in feeding nestlings after the incubation period; Turner & Rose 1994). Because prey size had no detectable relationship with flight performance (Table 3) and incubation type (Hasegawa & Arai 2022), it is also unlikely that foraging efficiency measured as prey size (sensu Turner 1982) indirectly affected the relationship between male incubation and flight performance (e.g., through differential time budget for foraging and paternal care). Rather, limited or frequent opportunities for extrapair mating in biparental or female-only incubation would explain the observed pattern, perhaps through female sire choice favoring fast and maneuverable flyers, through sperm competition for rapid and efficient sneaking, successful mate guarding, or combinations of these factors. Evolutionary transition analysis reinforces this perspective, because biparental incubation (i.e., limited opportunities for extrapair mating) and non-straight flight are evolutionarily incompatible (Fig. 4).

Some flight performance is known to evolve with sexual selection, particularly when used as courtship display (e.g., aerial display found in polygynous birds; e.g., Mikula et al. 2022). It is thus not surprising that our measures of flight performance evolved with an index of opportunities of extrapair mating, even when we evaluated daily flight performance (see Methods). In fact, sexual selection continues after the courtship period (i.e., extrapair mating, in this case; Møller 1994; Turner 2006; Romano et al. 2017), and it is straightforward to see that male traits that directly affect paternity can be targets of sexual selection (e.g., intensity of mate guarding: Møller 1987; also see Crino et al. 2017 for paternity success of male zebra finches with superior flight performance). However, in many empirical studies, sexual selection is often ignored in the context of evolution of locomotor performance and its interrelationship with ornamentation: few studies have attempted to link locomotor performance with indices of sexual selection (reviewed in Husak & Fox 2008; but see Husak et al. 2006, 2008; Chrino et al. 2017 for some exceptions). As a consequence, performance function of ornamentation is often thought to reflect the cost of ornamentation under the assumption that high performance, even if it accompanies viability disadvantages in unmeasured aspects, is favored by viability selection (e.g., Buchanan & Evans 2000; also see Brown & Brown 2013 for viability disadvantage of landing swallows with specific wing morphologies). Future studies should carefully take into account the potential importance of sexual selection on locomotor performance.

A caveat in the current study is that our measures of flight performance might be somewhat subjective to observers. For example, conspicuous performance (e.g., tight turn) would be more likely to be described than trivial performance, and thus, the former can be overrepresented in the current data. To minimize this possibility, we included any notable non-straight flight, such as zigzagging and erratic flight (i.e., we did not exclude “trivial” non-straight flight), but this might not be enough. The subjectivity is also applicable to fast flight, because whether or not birds are perceived to be fast would depend on their body size (i.e., small objects are perceived as “fast”), and possibly on their foraging habits (e.g., birds may be perceived as “fast” when they fly close to the ground). Thus, we should keep in mind that our measures of flight performance are based on how they are perceived, which might explain why opportunities for extrapair mating, a measure of intraspecific sexual interactions (see above), was related to flight performance in birds that share a part of visual systems with ours (e.g., Martin 2016). Objective measures of flight performance (e.g., flight velocity, turning angle, gliding duration) in each species are urgently needed.

In summary, we found here a quadratic (convex) relationship between notable non-straight flight and male tail fork depth in hirundines. Because incubation type, an index of opportunities for extrapair mating in hirundines, explains non-straight flight, flight performance and its link with tail fork depth can be affected by sexual selection at least in part. This can explain the apparent inconsistencies between theorical studies that propose an aerodynamic function for forked tails (see Introduction) and a series of macroevolutionary studies that do not support a viability function of forked tails (and instead support sexual function; e.g., Hasegawa & Arai 2020, 2021, 2022). A phenotype that provides the best performance can be away from the viability optimum, which then provides room for sexually selected, elaborate performance. The determination of phenotypes that provide the best performance would thus not clarify viability optimum ornament expression, preventing the inference of the evolutionary driver of ornamentation.

## Acknowledgments

I thank Dr. Emi Arai, Dr. Shumpei Kitamura and his lab members at Ishikawa Prefectural University for their kindest advices and supports. I also thank English-proofing company (Enago) for their helpful comments. MH was supported by the KAKENHI grant of the Japan Society for the Promotion of Science (JSPS, 19K06850, 22J40066).

---

**Table S1.**
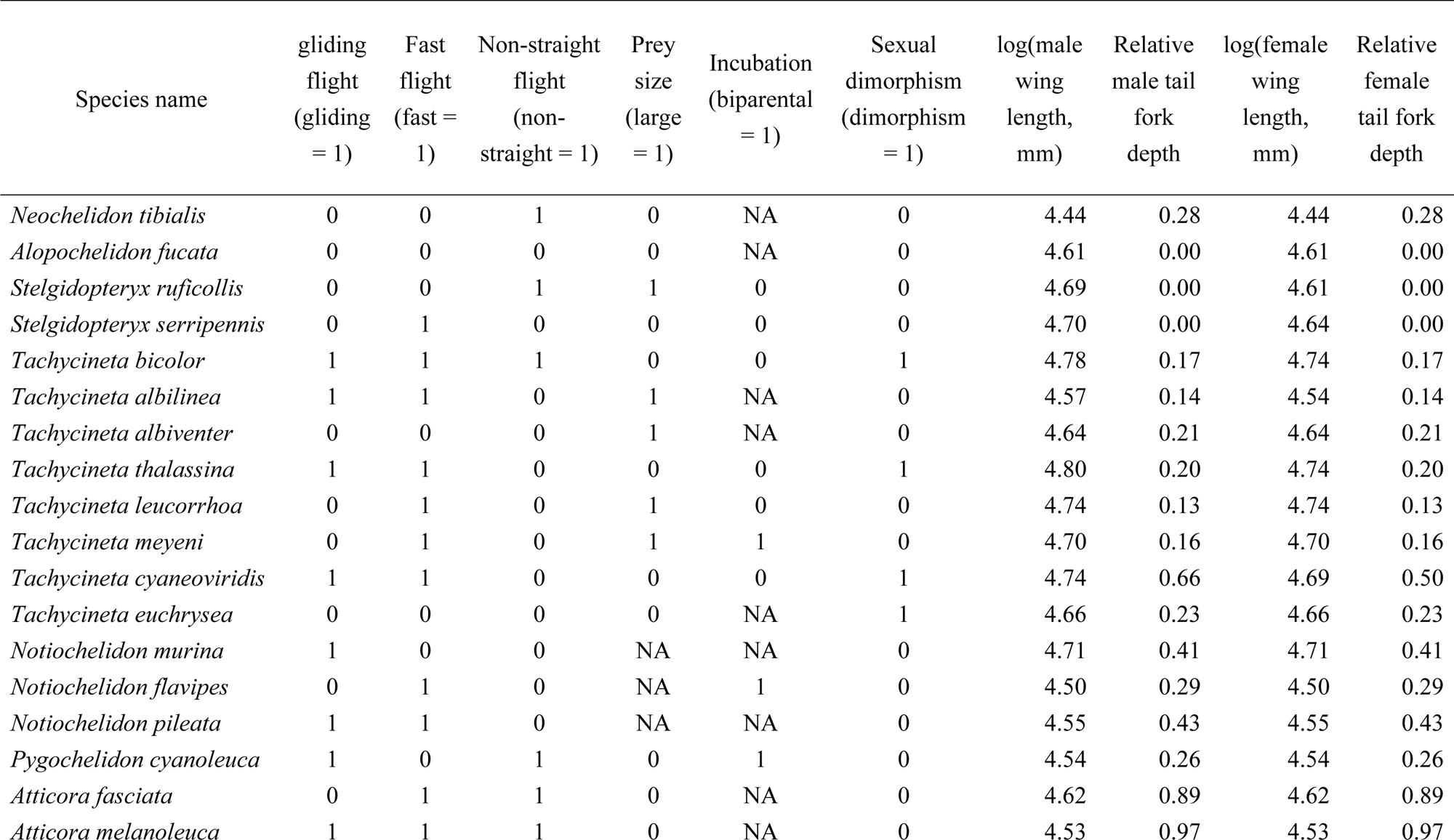
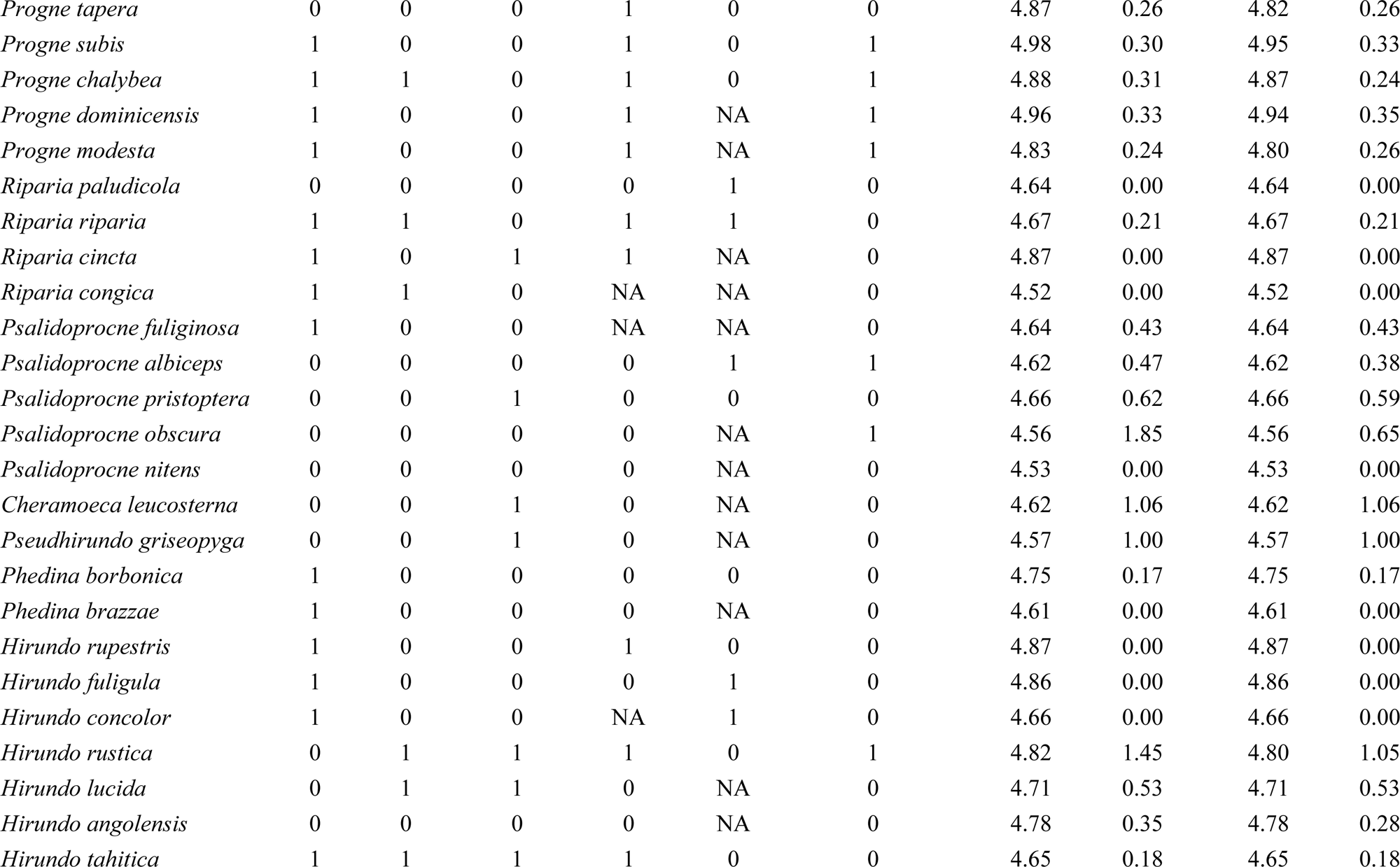
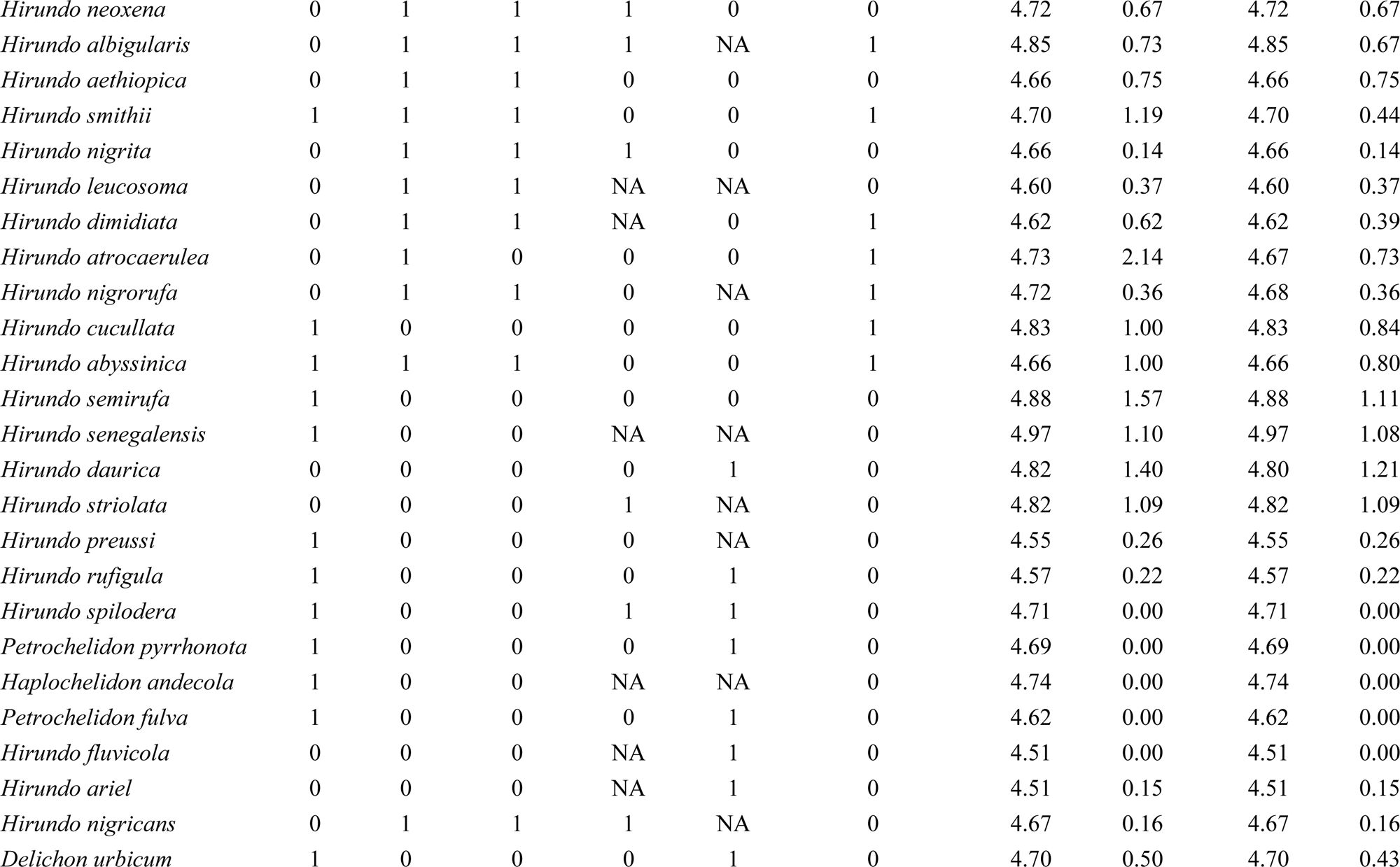
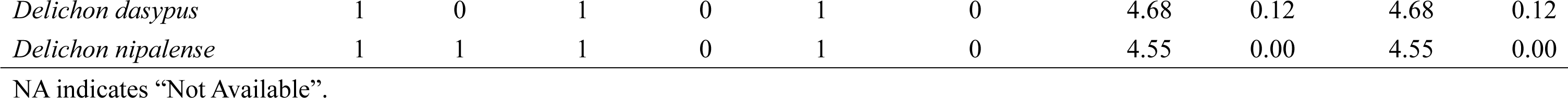
Dataset for the current study (n = 70).

**Table S2.**
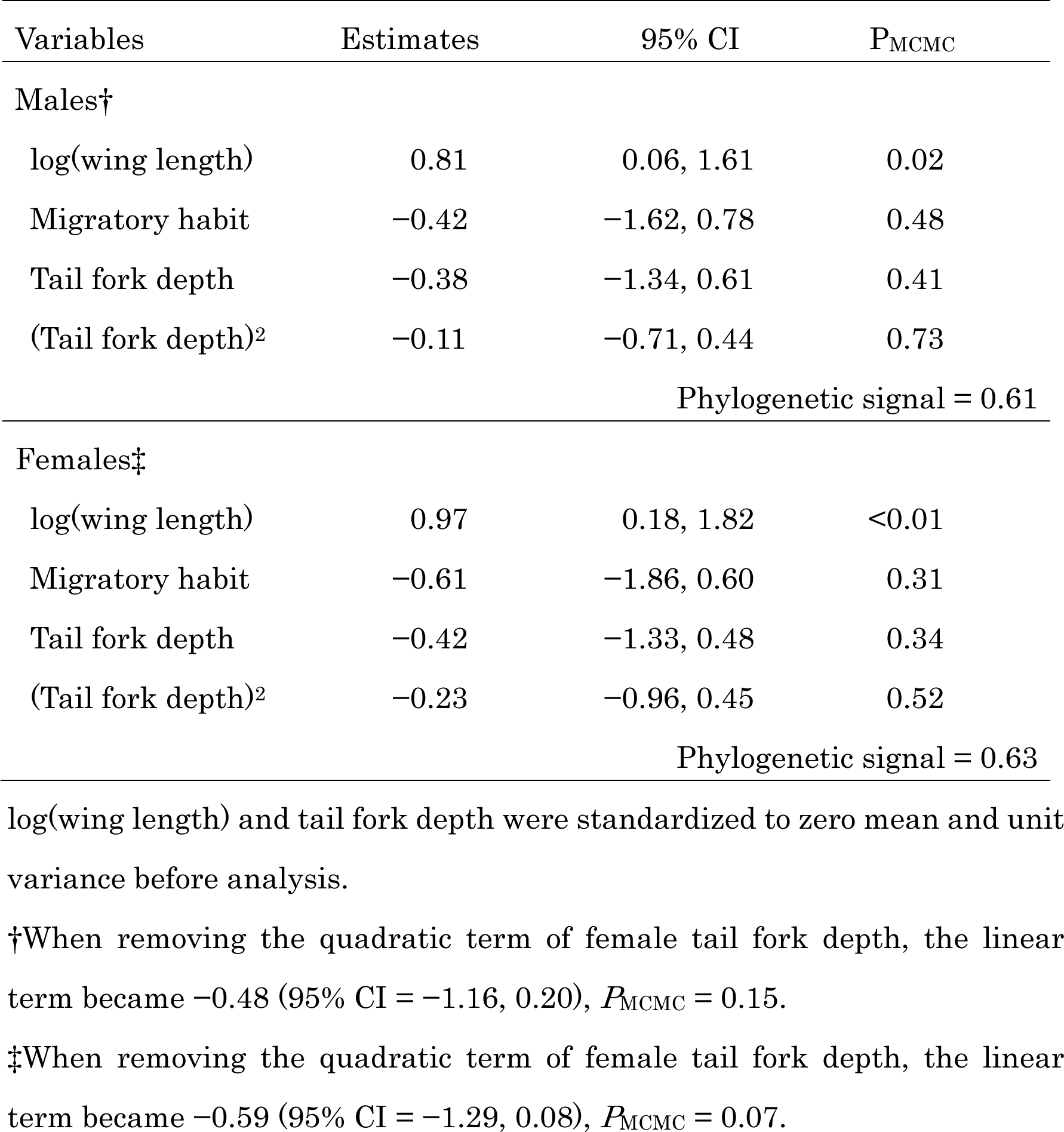
Multivariable Bayesian phylogenetic mixed model with binary error distribution predicting the presence or absence of gliding flight in relation to tail fork depth in males and females (both: n = 70).

| Variables | Estimates | 95% CI | P<sub>MCMC</sub> |
| --- | --- | --- | --- |
| Males† |  |  |  |
| log(wing length) | 0.81 | 0.06, 1.61 | 0.02 |
| Migratory habit | -0.42 | -1.62, 0.78 | 0.48 |
| Tail fork depth | -0.38 | -1.34, 0.61 | 0.41 |
| (Tail fork depth)<sup>2</sup> | -0.11 | -0.71, 0.44 | 0.73 |
| Phylogenetic signal = 0.61 |  |  |  |
| Females‡ |  |  |  |
| log(wing length) | 0.97 | 0.18, 1.82 | <0.01 |
| Migratory habit | -0.61 | -1.86, 0.60 | 0.31 |
| Tail fork depth | -0.42 | -1.33, 0.48 | 0.34 |
| (Tail fork depth)<sup>2</sup> | -0.23 | -0.96, 0.45 | 0.52 |
| Phylogenetic signal = 0.63 |  |  |  |

log(wing length) and tail fork depth were standardized to zero mean and unit variance before analysis.
†When removing the quadratic term of female tail fork depth, the linear term became -0.48 (95% CI = -1.16, 0.20), $P_{\text{MCMC}} = 0.15$ .
‡When removing the quadratic term of female tail fork depth, the linear term became -0.59 (95% CI = -1.29, 0.08), $P_{\text{MCMC}} = 0.07$ .

**Table S3.** Multivariable Bayesian phylogenetic mixed model with binary error distribution predicting the presence or absence of fast flight in relation to tail fork depth in male and female hirundines (both: n = 70).

| Variables | Estimates | 95% CI | P<sub>MCMC</sub> |
| --- | --- | --- | --- |
| Males† |  |  |  |
| log(wing length) | -0.77 | -1.63, 0.03 | 0.048 |
| Migratory habit | 0.34 | -0.95, 1.64 | 0.61 |
| Tail fork depth | 0.89 | -0.17, 2.00 | 0.09 |
| (Tail fork depth)<sup>2</sup> | -0.26 | -0.76, 0.26 | 0.31 |
| Phylogenetic signal = 0.69 |  |  |  |
| Females‡ |  |  |  |
| log(wing length) | -0.73 | -1.58, 0.06 | 0.056 |
| Migratory habit | 0.26 | -0.99, 1.54 | 0.69 |
| Tail fork depth | 0.96 | 0.01, 1.95 | 0.038 |
| (Tail fork depth)<sup>2</sup> | -0.61 | -1.32, 0.09 | 0.08 |
| Phylogenetic signal = 0.65 |  |  |  |

log(wing length) and tail fork depth were standardized to zero mean and unit variance before analysis.
†When removing the quadratic term of female tail fork depth, the linear term became 0.48 (95% CI = -0.23, 1.24), $P_{\text{MCMC}} = 0.18$ .
‡When removing the quadratic term of female tail fork depth, the linear term became 0.46 (95% CI = -0.27, 1.23), $P_{\text{MCMC}} = 0.20$ .

**Table S4.** Multivariable Bayesian phylogenetic mixed model with a binary error distribution predicting the presence or absence of gliding flight (upper column) and fast flight (lower column) in relation to incubation type as a measure of opportunity of extrapair paternity in male hirundines (both: n = 41).

| Variables | Estimates | 95% CI | P<sub>MCMC</sub> |
| --- | --- | --- | --- |
| Gliding flight |  |  |  |
| log(wing length) | 1.07 | 0.06–2.12 | 0.028 |
| Migratory habit | 0.17 | –1.50, 1.90 | 0.86 |
| Tail fork depth | –0.22 | –1.58, 1.13 | 0.75 |
| (Tail fork depth)<sup>2</sup> | –0.42 | –1.30, 0.38 | 0.32 |
| Incubation type | 0.64 | –1.14, 2.37 | 0.45 |
| Phylogenetic signal = 0.56 |  |  |  |
| Fast flight |  |  |  |
| log(wing length) | –1.11 | –2.27, –0.02 | 0.03 |
| Migratory habit | –0.86 | –2.37, 0.99 | 0.35 |
| Tail fork depth | 0.69 | –0.83, 2.26 | 0.37 |
| (Tail fork depth)<sup>2</sup> | 0.04 | –0.74, 0.89 | 0.96 |
| Incubation type | –2.20 | –4.14, –0.35 | 0.02 |
| Phylogenetic signal = 0.64 |  |  |  |

log(wing length) and tail fork depth were standardized to zero mean and unit variance before analysis.

**Fig. S1.**
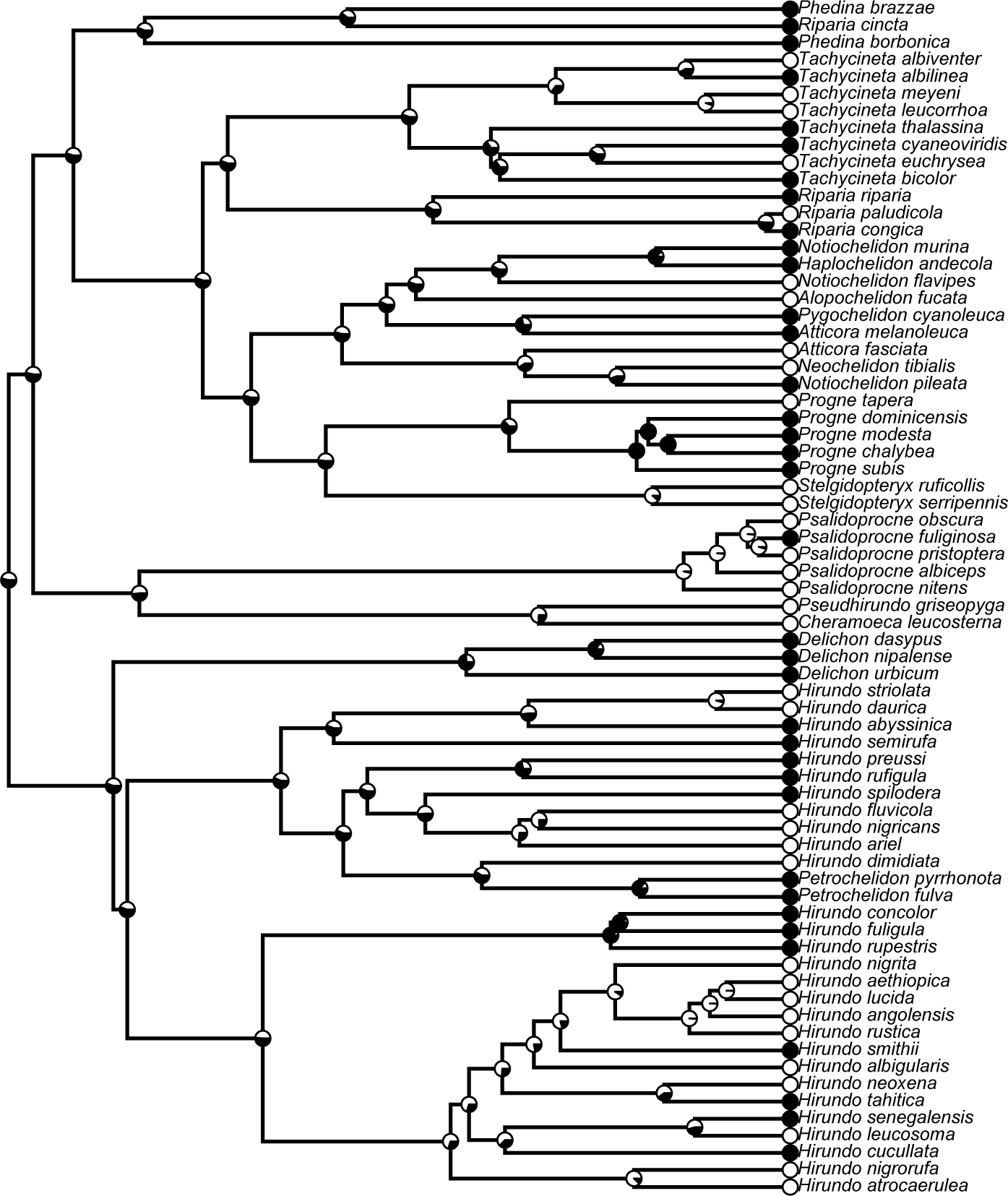
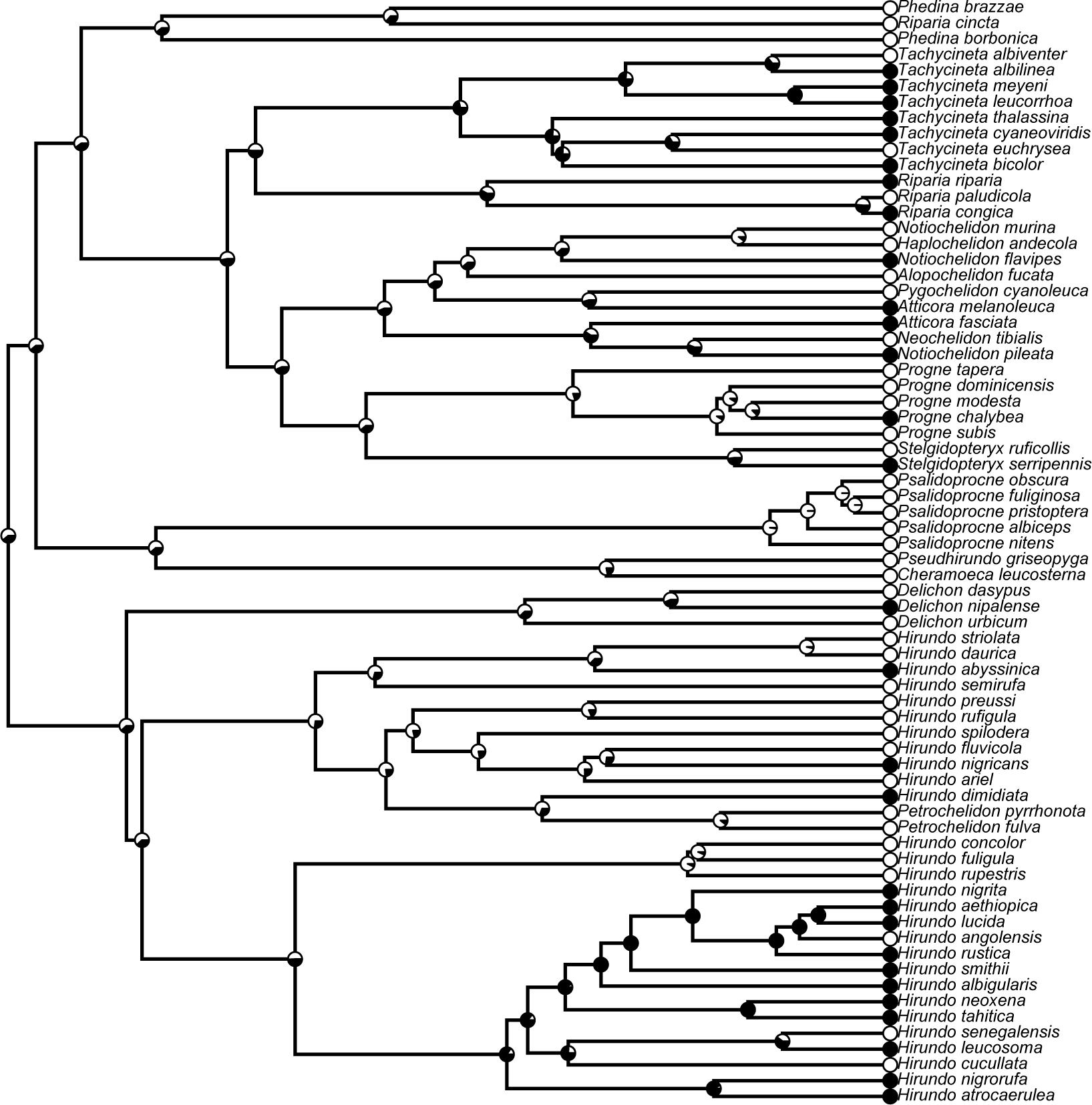
Examples of ancestral character reconstruction of gliding flight (this page) and fast flight (next page) in swallows and martins (Aves: Hirundininae). The presence and absence of each flight (i.e., gliding or fast flight) are indicated with black and white circles, respectively. The proportions of black and white at the nodes indicate the probability of the ancestral state. Here, I used the “ace” function in the R package “ape” (with model = “ER,” i.e., an equal-rates model; Paradis et al. 2005) and “plotTree” in the R package “phytools” (Revell 2012) for reconstructing ancestral character.

## Notes

### Competing Interest Statement

The authors have declared no competing interest.

